# Ruling the unruly: Innovation in ant larval feeding led to increased caste dimorphism and social complexity

**DOI:** 10.1101/2022.12.08.519655

**Authors:** Arthur Matte, Adria C. LeBoeuf

## Abstract

Building differences between genetically equivalent units is a fundamental challenge for every (super)organism with reproductive division of labor. In ants, reproductive or worker fate is typically determined during the larval stage. However, the methods by which adults feed their larvae, thus controlling their development, vary widely across ant species. Similarly, the body size gap between queen and worker is highly heterogeneous, ranging from species with similar-sized individuals to species with queens over 300 times larger than their smallest workers. To investigate the role of alloparental feeding control in caste dimorphism and the evolution of social complexity, we assembled data for queen:worker dimorphism, alloparental care, and larval morphology for a phylogenetically comprehensive sample of several hundred species, along with ecological and life-history traits. Using comparative phylogenetic methods, we analyzed the macroevolution of ant larvae and queen:worker dimorphism on a large scale. Our findings indicate that both extended alloparental feeding care and dimorphism are associated with the evolution of passive larval morphologies. Furthermore, greater queen:worker dimorphism co-evolved with several traits indicative of social complexity, including larger colony sizes, distinct worker subcastes, and the loss of full reproductive potential in workers. In sum, change in larval feeding habits were promoted by dietary shifts from prey to foods necessitating individualized distribution. These innovations granted adults greater capacity to manipulate larval nutrition, and consequently, caste size inequality, with significant implications for social complexity.

**Significance statement:** Ants are among the rare organisms to have extended reproductive division labor beyond the cells of a multicellular organism. However, the degree of specialization between reproductive and worker castes varies considerably between ant lineages. In this study, we demonstrate that strong caste dimorphism in ants co-evolved with complex eusociality traits, and this strong caste dimorphism was achieved by asserting adult control over larvae’ development. We conclude that this enhanced control over larval caste fate was a critical junction in the major evolutionary transition of ants toward caste specialization.

**Graphical abstract:** 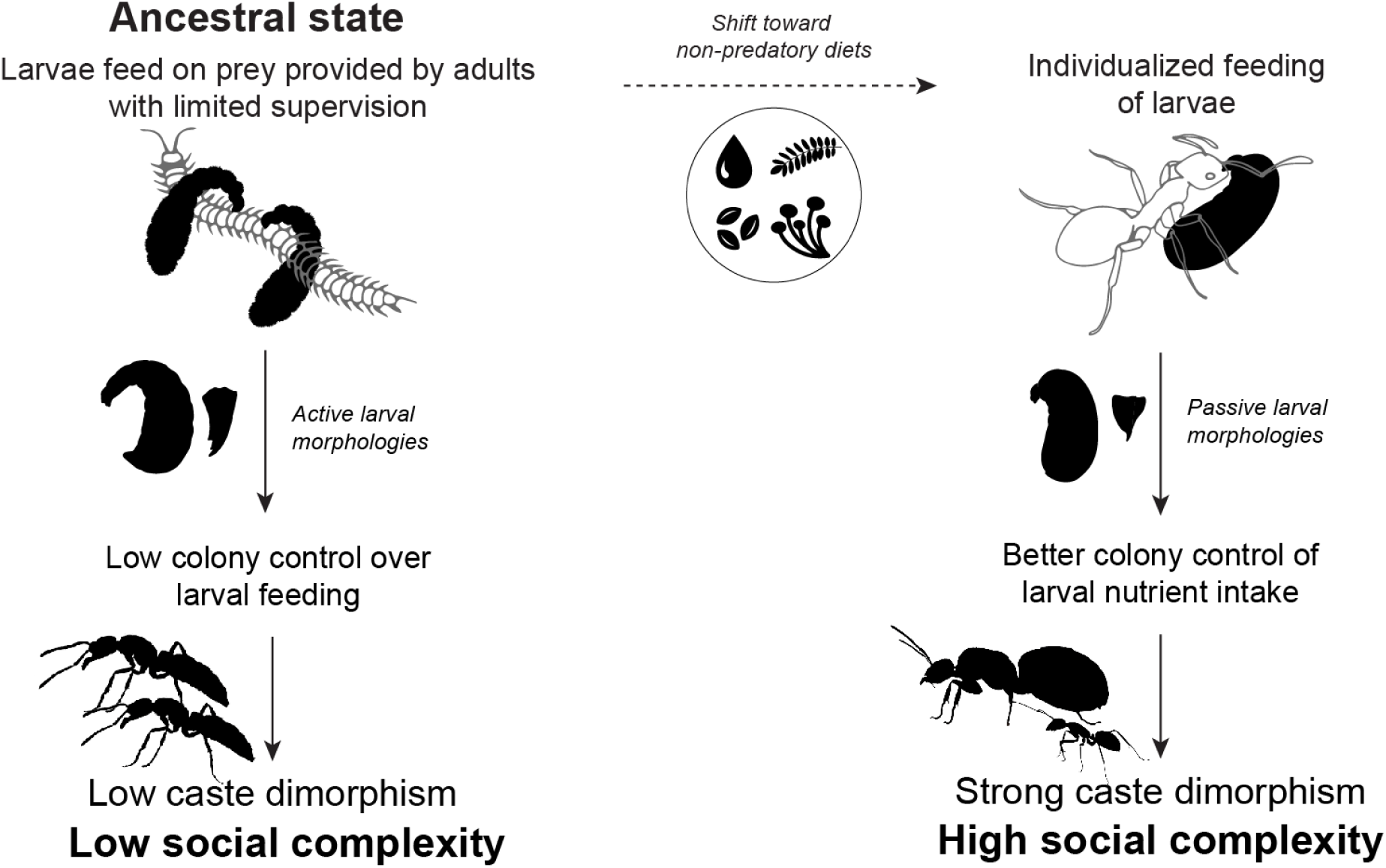

## INTRODUCTION

From the specialization of germline and somatic tissue in multicellular organisms to the sterile workers and reproductive queens of social insect colonies, reproductive division of labor is a central feature of deep cooperative systems (West et al. 2015, Boomsma and Gawne 2018, Holland and Bloch 2020). In eusocial Hymenoptera, adults are divided into reproductive and nonreproductive castes (Boomsma and Gawne 2018). Queens specialize in reproduction while workers specialize in food collection and brood care.

In ants, the size difference between queens and workers reflects the degree of caste specialization into their respective tasks (Trible and Kronauer 2017). Higher caste dimorphism is associated with bigger colony sizes (Ohyama et al. 2023) and benefits dispersing queens during colony foundation where smaller workers are comparatively cheaper to produce (Peeters 2020). Despite the purported advantages of higher caste dimorphism for social life, the morphological gap between queen and worker castes is highly heterogeneous across the ant phylogeny. At one extreme, early-branching lineages often have small colonies of predatory specialists with minimal morphological differences between reproductives and workers. At the other extreme, speciose ecosystem-dominating generalist species tend to have colossal colonies and high levels of queen:worker dimorphism (Peeters 1997, Keller and Peeters 2020). To our knowledge, no broad comparative studies have yet been conducted on caste determination in ants, leaving unclear the evolution of dimorphism and its consequences on ant evolution.

The larval stage is critical in building caste dimorphism. Female embryos of social Hymenoptera are generally totipotent and can turn into either queens or workers. While caste fate is specified as early as in the egg in a few species (Anderson et al. 2008, Weyna et al. 2021, Qiu et al. 2022, Schultner et al. 2023), the prevailing belief is that ’environmental’ (social) caste determination occurs during larval development for most species (Anderson et al. 2008, Schwander et al. 2010). In holometabolous insects, building size-differentiated castes relies on a non-homogenous distribution of larval feeding (Mirth and Riddiford 2007). Beyond size, these feeding differences can activate or repress different biochemical and gene expression pathways (Wheeler 1986, Rajakumar et al. 2018, Qiu et al. 2022), also influenced by hormonal supplies such as juvenile hormone (Wheeler and Nijhout 1981, Negroni and LeBoeuf 2023a).

For optimal cooperation at the colony scale, larval caste fate – and therefore the feeding of larvae – should be managed by workers rather than the larvae themselves. Larval totipotency brings about an intense conflict between each larva and the rest of the colony. Specifically, larvae would benefit by becoming queens rather than workers, thereby gaining greater direct reproduction, whereas the colony more often benefits more by producing workers (Bourke and Ratnieks 1999, Ratnieks and Wenseleers 2008, Ratnieks and Helanterä 2009). In cases where caste is likely to be self-determined, as in Melipona bees, many females selfishly choose to become queens at the expense of colony productivity (Wenseleers et al. 2003, 2004, Oliveira et al. 2022, Ferreira et al. 2024). In some stingless bees, larvae can even break into neighboring cells to obtain additional food and consequently develop into queens (Faustino et al. 2002, Ribeiro et al. 2006). Nutritional caste control can be highly effective in preventing excess queen production (Wenseleers et al. 2003). Honeybees employ size-delimited cells to rear worker-destined larvae, preventing unchecked growth. However, if honeybee larvae are reared outside of the colony, they exhibit a wide range of intermediates between workers and queens (Linksvayer et al. 2011, Leimar et al. 2012). In ants, where larvae are not enclosed, such control can only be exerted through alloparental care.

Although alloparental care toward larvae was likely a key feature of early ant societies (Hanna and Abouheif 2021, Rees-Baylis et al. 2024), the extent of feeding care varies widely among ant lineages. Larvae of some species are exceptionally autonomous, feeding directly on entire prey retrieved by adults (e.g., Masuko 1990). Whereas other species exhibit completely passive larvae fed by nurse workers which regurgitate liquids rich in endogenously produced components (LeBoeuf et al. 2016, 2018) in regular and time-calibrated one-to-one interactions (e.g. Cassill and Tschinkel 1995). Semi-autonomous larvae exhibit active feeding behavior and several authors have hypothesized that this less-supervised feeding method leaves more room for the reproductive aspirations of larvae, limiting the possibility to build dimorphism in these species (e.g., Wheeler 1900, Rüger et al. 2008, Penick and Liebig 2012). In those species, adults sometimes even aggress their larvae to prevent them from becoming queens (Brian 1973, Penick and Liebig 2012), though developmental mechanisms remain unclear.

In this macroevolution study, we combined data on morphology, ecology, life history traits and behavior across hundreds of species to unravel the evolution of caste size dimorphism in ants. Our findings indicate that both extended alloparental feeding care and dimorphism are associated with the evolution of passive larval morphologies. In turn, greater queen:worker dimorphism co-evolved with several traits indicative of social complexity, including larger colony sizes, distinct worker subcastes, and the loss of full reproductive potential in workers. We further investigated how dietary and behavioral shifts accompanied key innovations to the feeding care of adults towards their larvae. Overall, our study analyzes the traits that have driven and being the consequences of the development of pronounced size asymmetries between queens and workers.

## RESULTS

To investigate and understand the role of alloparental care in building caste dimorphism, we first established larval morphology as a proxy for alloparental care analyzing the evolution of larval morphology and its link with larval feeding habits and diet. Next, we analyzed the evolution of caste dimorphism and tested its correlation with larval morphology and other traits underpinning advanced sociality.

### Alloparental care, larval morphology, and diet evolution

Because details about alloparental care and larval feeding are scarce in the literature (Meurville and LeBoeuf 2021), we used larval morphological features as proxies for larval feeding habits. Ant larval morphology is exceptionally diverse (Wheeler and Wheeler 1976, Fig. 1) and is known to provide a signature of larval feeding methods (e.g., Wheeler 1918, Petralia and Vinson 1979, Masuko 2008). George C. Wheeler and Jeanette Wheeler underscored, throughout their extensive 66-year study on ant larvae, that passive larval morphological features correlate with greater adult assistance in larval feeding. Specifically, the lack of an elongated neck indicates larvae with reduced capacity for self-directed feeding, and instead, passive reception of food provided by adult workers. Second, the poor development of larval mandibles indicates feeding on soft food (e.g., on liquid passed by trophallaxis or on eggs), while larvae with medial teeth on their mandibles likely chew through tougher food (Wheeler and Wheeler 1951, 1976, 1986a).

**Figure 1.**
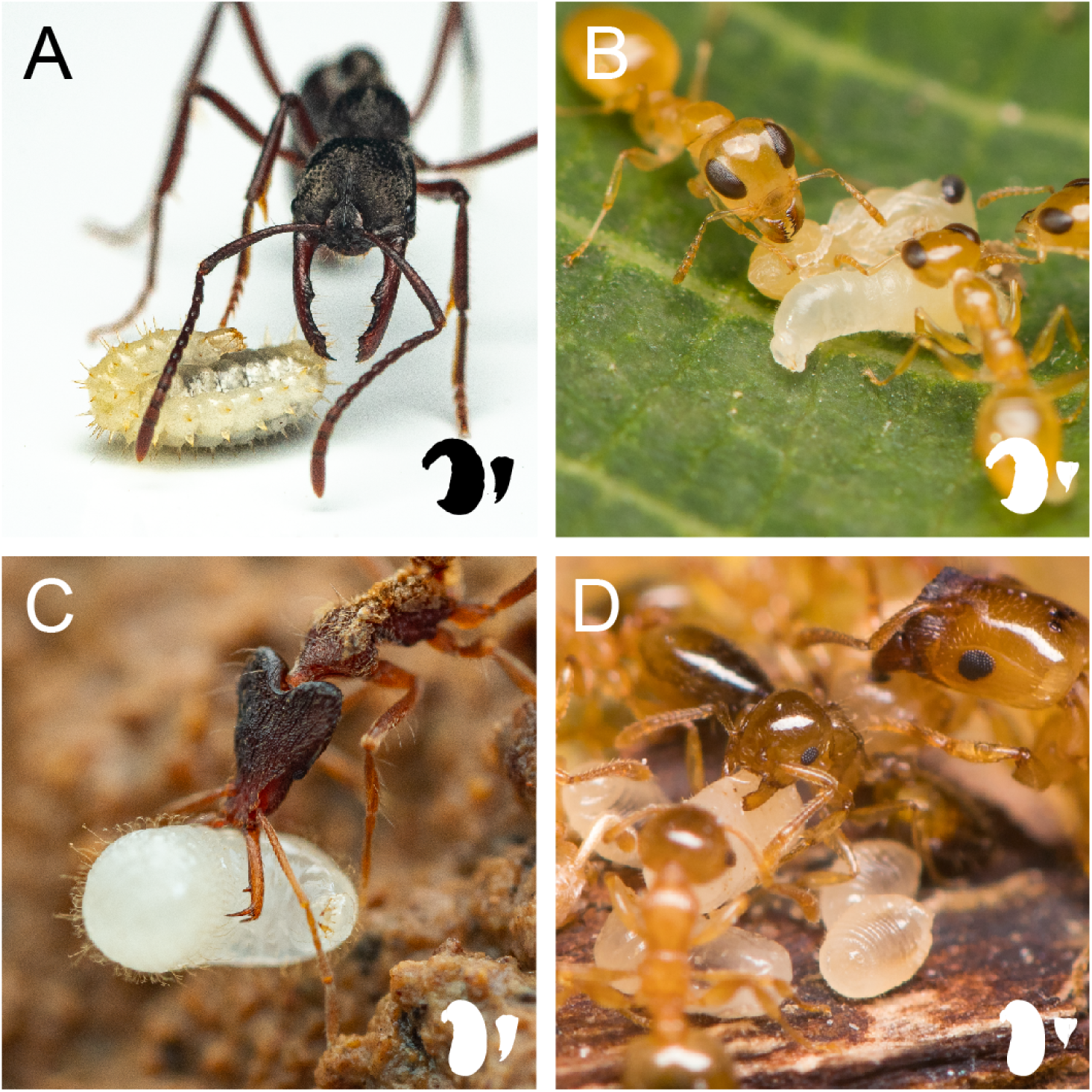
A sample of the diversity of ant larval morphology. **A)** *Myopias sp*: larva with elongated neck and developed mouthparts. **B**) *Gesomyrmex howardi*: larva with elongated neck and reduced mouthparts. **C**) *Strumigenys sp*: larva with reduced neck and developed mouthparts. **D**) *Crematogaster cylindriceps*: larvae with reduced neck and reduced mouthparts. Photos by François Brassard and used with permission.

We assembled data for the presence of an elongated neck and medial teeth in the larvae of 733 species to analyze the evolution of these two key morphological features. We tested whether these larval features indeed correlated with larval feeding habits using Pagel’s (1994) method on a set of 99 species spanning 71 genera where both behavior and morphology were available (Figure S1). Specifically, we categorized whether larvae of these species usually fed semi-autonomously with limited alloparental care (*viz*., actively feeding upon unprepared prey left at their disposal or receiving large chunks of food items) or, conversely, if their feeding primarily involves strong adult care (*viz*., regurgitation, preparation of pre-masticated pellets, distribution of trophic eggs, etc.) (see Supplementary material 1 {data feeding pattern} for quotes and classifications). We found a significant positive correlation between the evolution of neckless and toothless larvae and the evolution of more supervised alloparental feeding (both *p* < 0.001, Table S1). Additionally, we found supervised larval feeding habits significantly correlated with non-predatory adult diets (*p* < 0.001, Table S1).

To clarify the evolutionary relationships between adult dietary shifts, larval feeding habits, and larval morphologies, we employed phylogenetic path analysis to assess which of five plausible evolutionary models is best supported by the distribution of these traits over the ant phylogeny (Fig. S2A). One of the five models was significantly better than the others (Fig. S2B, illustrated in Fig. 2). This model indicates that when adult ants evolved a non-predatory diet, larval feeding habits shifted to less independent feeding manners, and in turn, this non-autonomous feeding led to shifts in larval morphology. Specifically, non-autonomous feeding was associated with passive larval morphologies (i.e., neckless and toothless).

**Figure 2.**
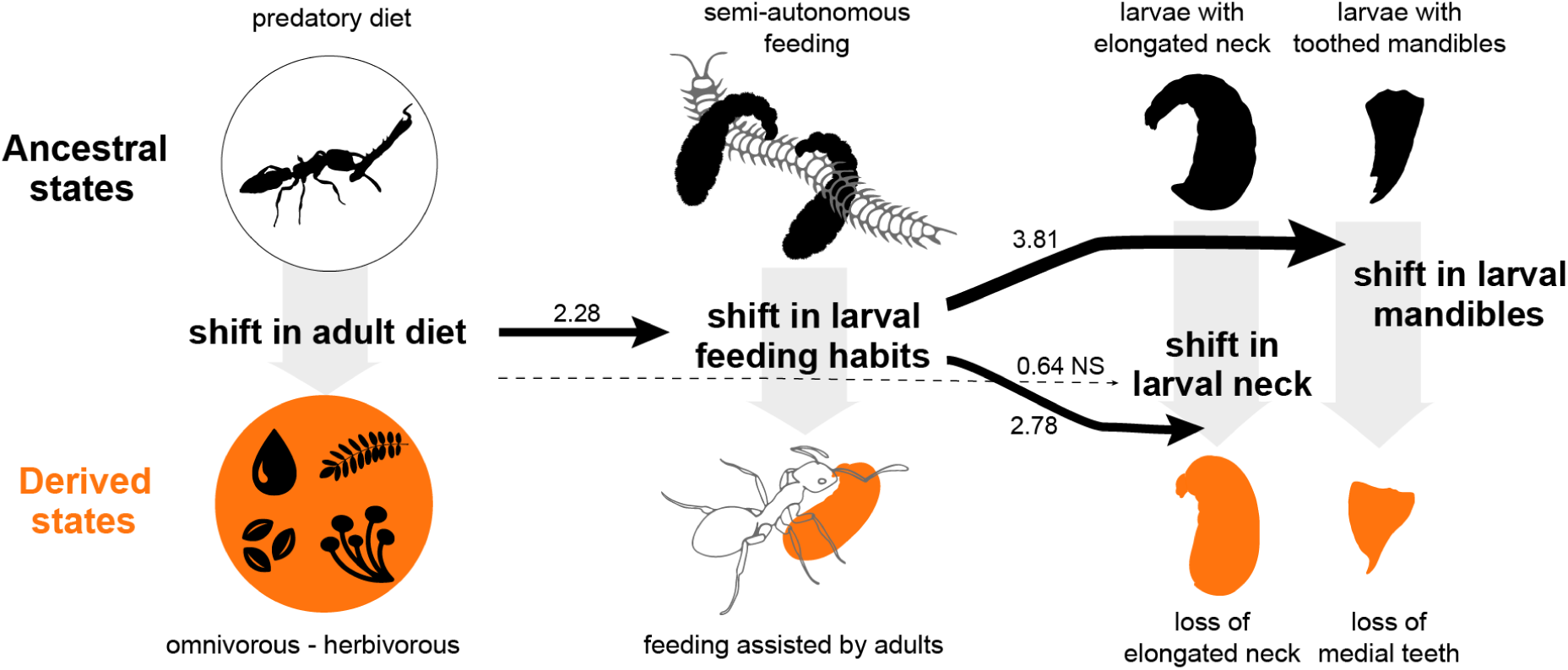
Trait relationships between diet, larval feeding habits and larval morphologies. Phylogenetic path analysis performed on a dataset of 99 species (71 genera), showing causal relationships between the studied variables from best supported model selection. Line thickness indicates the phylogenetic path coefficients and the dashed line indicates non-significant interactions (i.e., 95% CI including 0; see Fig. S2C). Ancestral diet state is supported by Smith et al. (2023) and ancestral larval feeding habits and morphology are supported by Fig. 3.

### Evolution patterns of larval morphology

We performed ancestral state reconstructions of larval morphology (neckless-toothless, necked-toothless, neckless-toothed, necked-toothed), according to three models with different constraints in discrete characters state changes and selected the best model upon the Akaike information criterion weight (AICw). The symmetric model was the best fit (AICw = 0.75, symmetrical transitions for each trait) compared with all-rates-differ (AICw = 0.22) and equal-rates (AICw = 0.02, all transition rates equal). The ancestral state reconstruction of larval morphology revealed multiple independent evolutions of neckless and toothless larvae from a necked-toothed ancestor (Fig. 3). The evolutionary pattern of acquisition of neckless and toothless morphology aligns with the Wheelers’ respective associations of these morphologies with an individualized distribution of food and specialization in soft foods (see above), and highlights the different pathways taken by ant lineages to feed their young.

**Figure 3.**
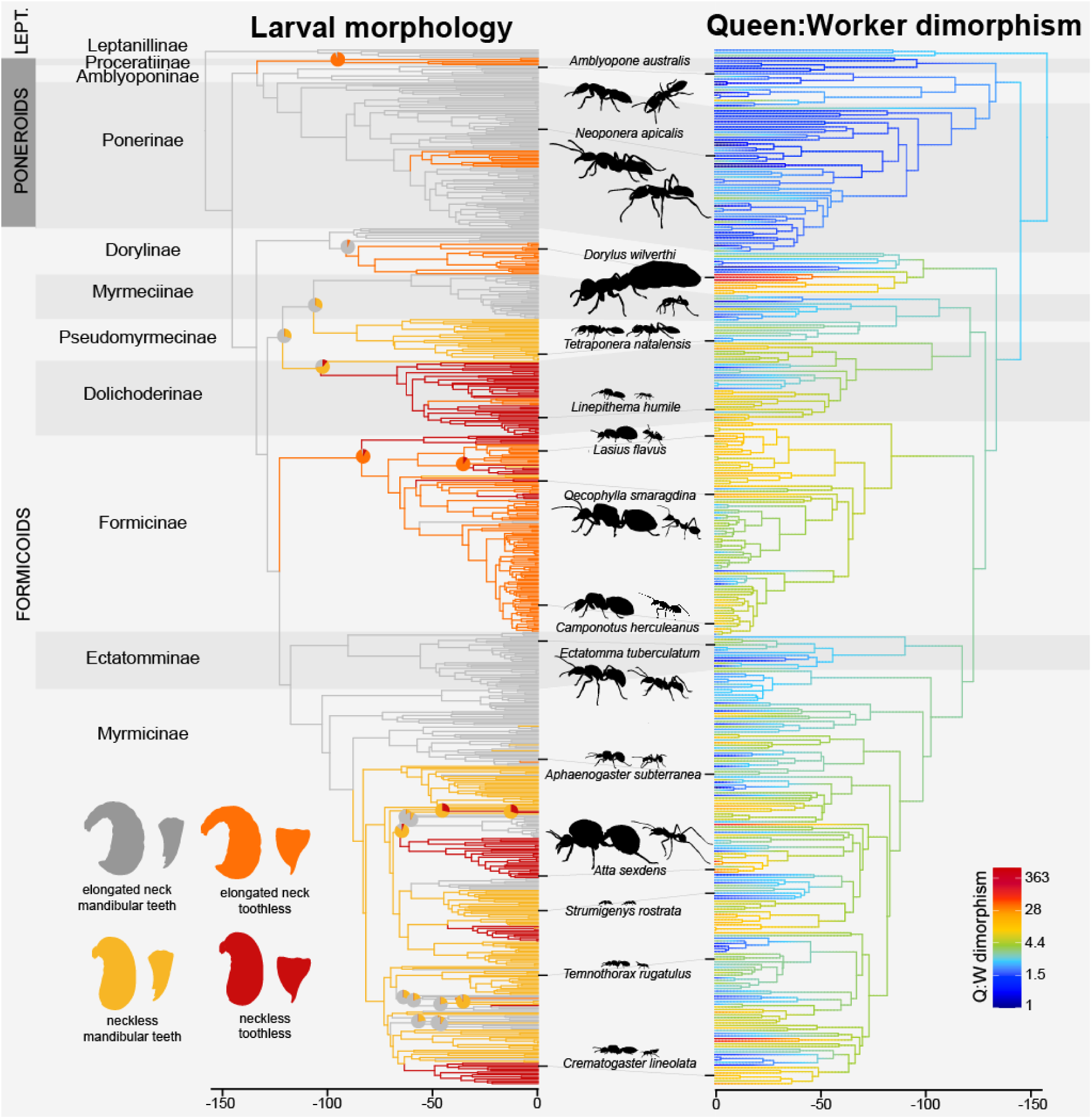
Ancestral state reconstructions of (left) larval morphology classified for 733 species and (right) queen:worker volume dimorphism measured on 392 species. Colored branches indicate predicted values of traits. Pie charts are plotted on nodes where the confidence < 95%. Silhouettes of queens (left) and workers (right) are shown in proportion to their measured size. Major subfamilies are labeled in alternating tones with details of their clade: e.g., “LEPT.” = Leptanillomorphs. Time is expressed in Ma and the color bar on the right-hand panel indicates the ratio between queen and worker body volumes.

Neckless larvae have predominantly evolved in Dolichoderinae, Pseudomyrmecinae, and Myrmicinae. Current evidence indicates that Dolichoderinae primarily nourish their larvae with trophic eggs (ref), while Pseudomyrmecinae larvae mostly feed on pellets prepared by adults (Wheeler and Bailey 1920). In lineages of Myrmicinae that acquired neckless larvae early in the subfamily’s evolution, this morphology is found in extant species exhibiting a generalist diet and feeding their larvae through regurgitation and pellets (e.g., *Solenopsis*, *Monomorium*, *Temnothorax*, *Pheidole*). However, larvae of some species regained an elongated neck, in particular in species exhibiting primarily predatory diets (e.g., *Orectognathus, Epopostruma, Eurhopalothrix, Myrmecina, Terataner*). Conversely, necklessness was followed by the subsequent loss of medial teeth in larvae of lineages that largely ceased feeding their larvae with prey (e.g., fungus-growing ants, *Cephalotes*, *Crematogaster*). Neckless larvae evolved in species exhibiting individualized distribution of food, regardless of the nature of the food.

Toothless larvae evolved in six subfamilies: at or near the root of the subfamilies Formicinae, Dolichoderinae, Proceratinae, and more sparsely in Myrmicinae, Ponerinae, and Dorylinae (Fig. 3). Aside from Dorylinae and Ponerinae, these ant species with toothless larvae are all documented to predominantly feed their larvae through trophallaxis, eggs, or fungal staphyla (Supplementary material 1 {data feeding pattern}), aligning with larval specialization on soft food. Within Formicinae, our dataset revealed that the only observed reversions to necked and toothed larvae occurred in *Myrmoteras* and possibly in *Proformica*. *Myrmoteras* has undergone dietary and behavioral reversion towards specialization on prey capture and displays phylogenetically uncharacteristic low dimorphism between queens and workers (Ito et al. 2017). Within Dorylinae, toothlessness evolved only in the army ants.

We hypothesized that these shifts in adult control over larval nutrition might have enabled the development of different adult female morphologies by allowing the colony’s workers to direct larval fate according to the colony’s needs and maturation. To test this hypothesis, we focused on the most fundamental female morphological difference in ants, between queens and workers. While queens and workers show a broad array of morphological differences, we simplified their differences by focusing only on volumetric body-size dimorphism.

### Evolution of queen:worker volume dimorphism

Ants exhibit a large range of queen:worker dimorphism within their evolution. Our taxon sampling includes a wide range of dimorphisms, ranging from species with similar sized castes (e.g., in *Myrmecia, Neoponera, Prionopelta*), to species having queens over one hundred times bigger than their smallest workers (e.g., in *Atta, Carebara, Dorylus, Tranopelta*). Phylogenetic analysis of the evolution of queen:worker dimorphism revealed a lambda model of evolution with a phylogenetic signal *λ* of 0.85 (Table S2), indicating that dimorphism is slightly less similar amongst species than expected given their phylogenetic relationships (Pagel 1999).

Despite considerable variability in ant queen:worker dimorphism, the Formicoids stand in stark contrast to the Poneroid and Leptanillomorphs clades (Fig. 3). This same branch in the ant phylogeny was previously found to have sustained rapid positive selection (Romiguier et al. 2022). Notably, 90% of sampled poneroid species and 100% of the sampled Leptanillomorphs exhibit low dimorphism (Q:W_volume_ < 3). In contrast, formicoid ants display extreme variation in dimorphism, with significantly higher average dimorphism values in Dorylinae, Myrmicinae, Formicinae, and Dolichoderinae than in Poneroids (one-way ANOVA: F_11,376_ = 15.812, p < 0.001; Tukey’s post hoc test: p < 0.05, Fig. S3).

These stark differences cannot be attributed to potential limits in the body volume evolution. Queen and worker body volumes shifted dramatically, both increasing and decreasing in all ant lineages (Fig. 3) with no clade-specific difference in evolutionary rates (Likelihood Ratio Test: both *p = 1*). Queen and worker body volume evolution were best fit with an Ornstein-Uhlenbeck and a delta model of evolution with phylogenetic signals *λ* of 0.87 and 0.89 respectively (Table S2). Queen volume evolved faster than did worker volume (σ^2^ *queen* = 9.3e10^-3^, σ^2^ *worker* = 5.8e10^-3^, Likelihood Ratio Test: *p* < 0.001) indicating putative different levels of selection pressure on the two castes.

### High queen:worker size dimorphism is associated with larval passiveness and traits of advanced eusociality

While body size of queens and workers can shift rapidly in all lineages, the difference between queen and worker size remained consistently low in some groups while variable in others. We next investigated the evolution of the size gap between these castes – queen:worker dimorphism. We found strong dimorphism predominantly evolved within the subfamilies exhibiting passive larvae (Fig. 3).

Formicoid lineages make up 80% of extant ant species and ants of these lineages are the most abundant ants worldwide (Borowiec et al. 2021). They have successfully colonized diverse ecological niches, often exhibiting generalist feeding behaviors, including both insect prey and nectar or honeydew consumption. In contrast, low dimorphism subfamilies (ants from the poneroid clade, but also Myrmeciinae and Ectatomminae) are primarily predators (Blanchard and Moreau 2017, Sosiak and Barden 2021).

Given these associations between the high dimorphism of most formicoid ants and other aspects of advanced eusociality, we wanted to assess whether queen:worker dimorphism was associated with larval passiveness traits (necklessness, toothlessness), more complex social organization and dietary shifts. We compiled a series of traits typically associated with advanced eusociality, including the absence of workers with full reproductive potential (e.g. gamergates, Monnin and Peeters 1999), colony size (Holbrook et al. 2011, Ferguson-Gow et al. 2014), and presence of polymorphic workers (Boomsma and Gawne 2018). We treated the number of passive traits in larval morphology as ordinal data (i.e., zero, one, or two). Additionally, we gathered dietary information, contrasting genera that are primarily predators with those that are omnivores or herbivores.

Our Phylogenetic Generalized Least Squares (PGLS) analysis revealed that larval passiveness (*p =* 1.9e10^-4^), larger colony size (*p* = 2.6e10^-5^), worker polymorphism (*p =* 1.9e10^-8^) were positively correlated with greater Q:W body size dimorphism. Workers with full reproductive potential were negatively correlated with Q:W body size dimorphism (*p* = 0.031). Predatory diet did not significantly correlate with Q:W body size dimorphism (*p* = 0.37). The detailed results of PGLS analyses including the relationship of these variables with queens and workers body volume are provided in Table S3.

To disentangle the relationships between our variables and their inherent causalities, we constructed four plausible evolutionary models (Fig. 4). These models were assessed and compared using a modified phylogenetic path analysis (van der Bijl 2018) to compute models and fit coefficients over multiple trees instead of a single one. Our models were designed to test for direct versus indirect associations between larval passiveness and Q:W dimorphism. The relationships common to all causal models are based on relationships supported by literature and also found to be significant in our PGLS analysis (Table S3). In addition to testing for a direct or indirect link between larval passiveness and dimorphism, we included models where changes in colony size led to changes in dimorphism and larval passiveness, and models where changes in dimorphism led to changes in colony size. The two best supported models were averaged (Fig. 4). Both the model choice and coefficients support a direct link between increased larval passiveness and increased Q:W dimorphism (Fig. 4).

**Figure 4.**
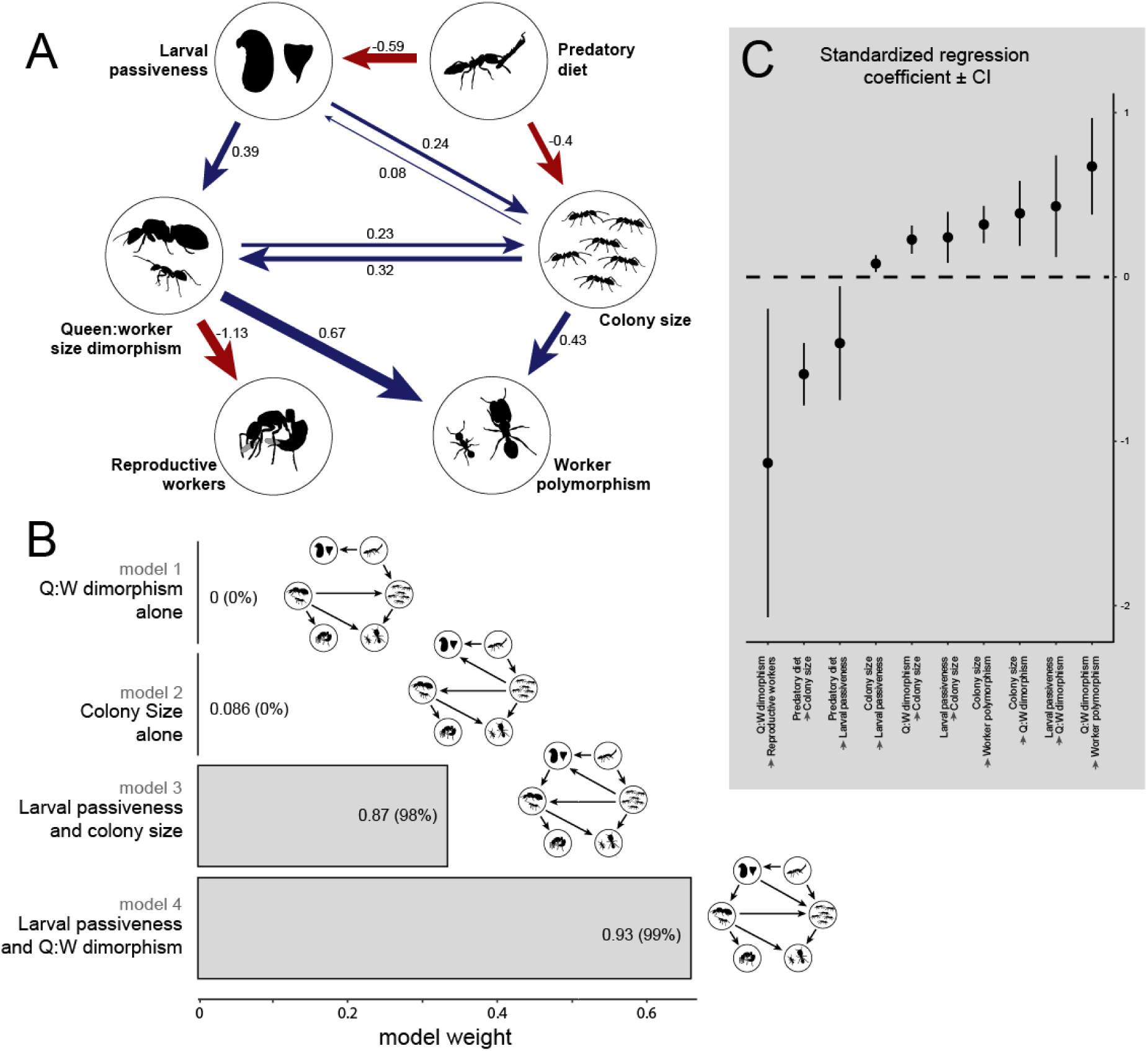
Examination of the causal relationships between larval passiveness, Q:W size dimorphism, colony size and other traits of advanced eusociality and ecology. Multi-tree phylogenetic path analysis performed on a dataset of 363 species (128 genera). **A**) causal relationships between variables from averaging the best models. Arrow values indicate the phylogenetic path coefficients. Blue edges indicate positive correlations while red edges mark negative correlations. **B**) Model comparison. Bar labels are p−values, significance indicates model rejection; values between parentheses are the percentages of model selection over 200 phylogenetic trees. **C**) standardized regression coefficient ± CI estimated over 200 phylogenetic trees from the selected models (models 3 and 4).

## DISCUSSION

Early ants were already capable of creating distinct queen and worker morphologies (Hanna and Abouheif 2021, Boudinot et al. 2022). Our study suggests that species became able to considerably increase their caste dimorphism, and reap considerable fitness benefits, by obtaining greater control over larval feeding. We found that low queen:worker dimorphism and actively feeding larvae were the ancestral states for ants. Dimorphism increased dramatically in several formicoid lineages, where it co-evolved with larval passiveness, large colony sizes, worker polymorphism, and loss of worker reproductive potential. The shifts of larvae from active feeders to passive food receptacles were promoted by pre-innovations shifting adult diets toward new food sources that required individualized distribution. We propose that shifting the ancestral feeding pattern of larvae toward extended and individualized nutritional alloparental care was a key evolutionary innovation that allowed the evolution of extreme dimorphism.

### Passiveness allows but does not necessitate extreme dimorphism

Factors other than larval feeding influence queen:worker dimorphism. Notably, there are low-dimorphism species featuring passive larvae and high-dimorphism species with active larvae. Colony foundation and queen dispersal provide significant constraints on the evolution of dimorphism. During solitary colony foundation larger dispersing queens can produce more comparatively cheap workers to quickly achieve a minimally viable colony (Peeters 2020). However, other dispersal solutions exist that do not require high dimorphism (Hakala et al. 2019), or that remove limitations to queen volume (e.g., no need for flight in Army-ant Dorylinae).

Strong dimorphism in species without passive larval morphology can also be achieved through more rare methods of larval caste fate regulation. Some species have caste fate partially or fully determined by genetic factors, biasing diploid larval totipotency (Anderson et al. 2008, Weyna et al. 2021). In some ant taxa, adults mildly injure their larvae and feed on their hemolymph. This non-destructive cannibalism pilfers nutrients from larvae, and enables the creation of nutritional asymmetries that can result in size dimorphism (Masuko 1986). Larval hemolymph feeding has been found to be particularly frequent in *Leptanilla japonica* and *Myopopone castanea*, which both exhibit phylogenetically unexpected high dimorphisms (Masuko 1986, Ito 2010). This mechanism of creating dimorphism requires deeper investigation.

### Enforced altruism and the role of passiveness in the evolution of inequality

Ants are believed to have evolved from a parasitoid ecological background (Blaimer et al. 2023), which consistent with our ancestral state reconstructions, would have required active larval morphology for independent feeding on adult-acquired prey. While the amount of food received by autonomous larvae on a carcass can still be adjusted by workers by removing larvae from prey, building dimorphism this way would require strong and consistent social organization. Over the evolutionary history of ants, several lineages evolved increased nutritional alloparental care towards their larvae. Instead of feeding semi-autonomously on prey or large pieces of food, larvae came to passively receive food from adults, often in the form of regurgitated liquids or trophic eggs provided through one-to-one interactions. This enhanced parental care granted adults increased control over the nutrients consumed by each larva. Such a level of control was likely needed to manage the selfish reproductive aspirations of larvae and to create asymmetric nutritional states, channeling larvae into distinct adult morphologies (queen, worker, different worker subcastes, etc.).

Alongside behavioral adaptations, physiological adaptations underlie adults’ control over larval development. In numerous species, passive larvae are extensively fed through trophic eggs or regurgitated liquids whose components are produced by the adults themselves (Supplementary material 1 {data feeding pattern}, Negroni and LeBoeuf 2023b). Trophic eggs have been reported to influence caste determination (Genzoni et al. 2023) and regurgitated fluids have been found to contain larval growth regulators that adjust adult body size (LeBoeuf et al. 2016, 2018, Negroni and LeBoeuf 2023a), providing mechanisms of how workers can modulate larval development through personalized nutrition.

### Benefits of larval passiveness

Putting larval fate under adult control was a critical change point in the social transitions of ants. If alloparental care toward brood enabled early ant societies to build the first female castes (Hanna and Abouheif 2021, Rees-Baylis et al. 2024), individualizing adult control over larval nutrition allowed ants to build stronger caste dimorphism and take a major step in their sociality. The ability to precisely channel larvae into different castes enabled greater reproductive and worker cast specialization, along with subcaste of workers specialized for different tasks (Wilson 1980), increased colony size (Peeters and Ito 2015, Trible and Kronauer 2017), and produce massive dispersing queens able to found new independent colonies with no external food supply, reducing predation risk (Peeters 2020). These outcomes were made possible through stronger colony-level control over their larval development. Considering the major evolutionary transition from single-celled to multicellular organisms, a comparable shift in individual to collective priorities likely necessitated non-autonomous fate determination during development (West et al. 2015, Veit 2019). This may be a fruitful avenue for future research on the mechanisms that allowed the evolution of multicellularity.

### How could passive larvae have evolved and why don’t all ants exhibit passive larvae?

The evolution of passive larvae is somewhat surprising given ants’ parasitoid evolutionary origins (Blaimer et al. 2023) and individual fitness incentives. In a hypothetical early ant colony where larvae are fed by being placed near prey without individualized care, each diploid larva should aim to be the largest such that she could develop into a reproductive queen and ensure her own fitness gains. Thus, the most passive larva within a colony has little opportunity to become a reproductive queen transmitting passive larval genes. Instead, the most unruly larva should be favored (e.g., Trible et al. 2023). Furthermore, the additional care required by passive larvae puts a larger care burden on workers.

Conversely, when workers have gained the power to control larval fate, this allows the colony-level priorities to supersede the individual-level priorities. Given the benefits to the colony level of evolving worker strong control of larval fate, why don’t all ant species exhibit passive larvae? The fact that passive larvae have evolved multiple times in different ways (toothless, neckless) despite the aforementioned evolutionary barriers suggests the existence of innovations preventing larvae from obtaining extra meals despite an active morphology, notably through obligatory individualized feeding care.

Dietary shifts may have provided the pre-innovations that enabled individualized care. We found that passive larval morphologies were associated with more supervised feeding habits, which in turn were linked to dietary shifts away from a predatory diet. Throughout ant diversification, numerous species have transitioned from predatory to omnivorous or even fully herbivorous diets (Blanchard and Moreau 2017). While prey items can be shared among larvae in joint feasts or distributed into large pieces, many more derived diets are unshareable. Sugary liquids, trophic eggs, Beltian and Mullerian bodies, fungal staphyla, and some seeds, due to their material nature and small size, require adults to feed larvae individually and in small quantities through one-to-one interactions. Thus, whereas ancestral ant adults could drop large pieces of prey on or near active larvae and remove them at their discretion, shifts in adult diet necessitated incremental and individualized feeding of larvae. However, transitioning to alternative food sources while maintaining nutritional support for larval growth required physiological or ecological innovations, often achieved through symbiont acquisition or metabolic division of labor (see supplementary discussion).

Change for more supervised and individualized larval feeding habits had three crucial outcomes. First, it reduced the opportunity for larvae to overfeed so as to influence their caste fate (but see Peignier et al. 2019). Second, it dramatically reduced potential direct food competition among larvae. These are two mechanisms promoting active larval morphologies otherwise. Third, shifts for non-predatory diet likely changed the nutrient flows within the colony. While predatory species often rely on larvae to process their high-protein meals and redistribute nutrients to adults, the converse occurs in carbohydrate- or lipid-based diets (Went et al. 1972, Dussutour and Simpson 2009, 2012). Hense, shifts to obligate individual feeding granted adults greater control over larval nutrition and facilitated the evolution of morphologically passive larvae, less capable of manipulating their own nutrient intake. Both paved the way for great queen:worker dimorphism and its myriad benefits for ecological dominance.

## MATERIAL AND METHODS

### Caste body volumes and dimorphism

To examine the evolution of dimorphism and its relationship with a panel of ecological and life-history traits, we measured body size of ant queens and workers in 339 species spanning 138 genera over 14 subfamilies. Species were selected based on the presence of both queen and worker image (Antweb.org), available data for our traits of interest in the literature, phylogenetic data and optimally wide taxon sampling (Fig. S5).

Using the Antweb.org photo database, we measured one queen and one worker per species, favoring type specimens when possible (56% of our dataset are from type specimens). As our goal was to evaluate the maximum size difference between queens and workers, intermediate forms of workers such as soldiers were not measured. Ergatoid queens (i.e., wingless at adult emergence) were not measured either, except when they represented the only known reproductive form for the species (*viz.*, several Dorylinae and *Blepharidatta conops*; Peeters 2012). Due to the lack of pictures for both queen and worker of Leptanillinae species with phylogenetic data on Antweb.org, we measured queens and workers body volume for one *Leptanilla* and two *Protanilla* species from six specimens illustrated in Ogata et al. (1995) and Hsu et al. (2017). While ideally, we would have measured queen-worker pairs from the same colony or locality, the availability of imaged specimens would have significantly restricted our sample size. Therefore, we utilized the available images to sample the widest possible range of species. Consequently, our measures of dimorphism might be influenced by geographical effects on queens and workers from different localities (e.g., Wills et al. 2014, Brassard et al. 2020). However, this effect should be randomly distributed across our dataset and thus have a limited impact compared to the actual species effect on dimorphism variation.

We measured the length, width and height of the mesosoma as well as the length and width of the head and the gaster using ImageJ (Collins 2007) (v.1.53a) (Fig. 6). Because measurements are sensitive to tagma orientation, head height was impossible to properly measure on numerous mesosoma-pinned specimens. We therefore used the dataset of Sosiak and Barden (2021) to estimate the best proxy for this metric. Head length explained 95% of the head height variation (n = 299), allowing to approximate head height as 0.608 x head length. Gaster length and width were taken from measured ellipses that matched as closely as possible the shape of the gaster, balancing the gaster areas inside and outside the ellipse when required (Fig. 6). Body volumes and dimorphism were estimated as:

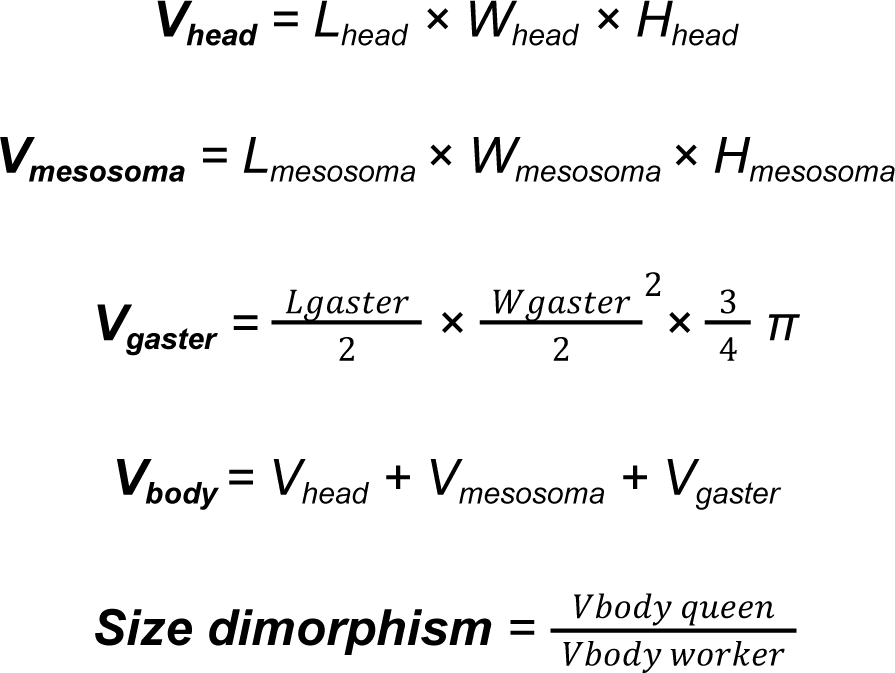

**Figure 6.**
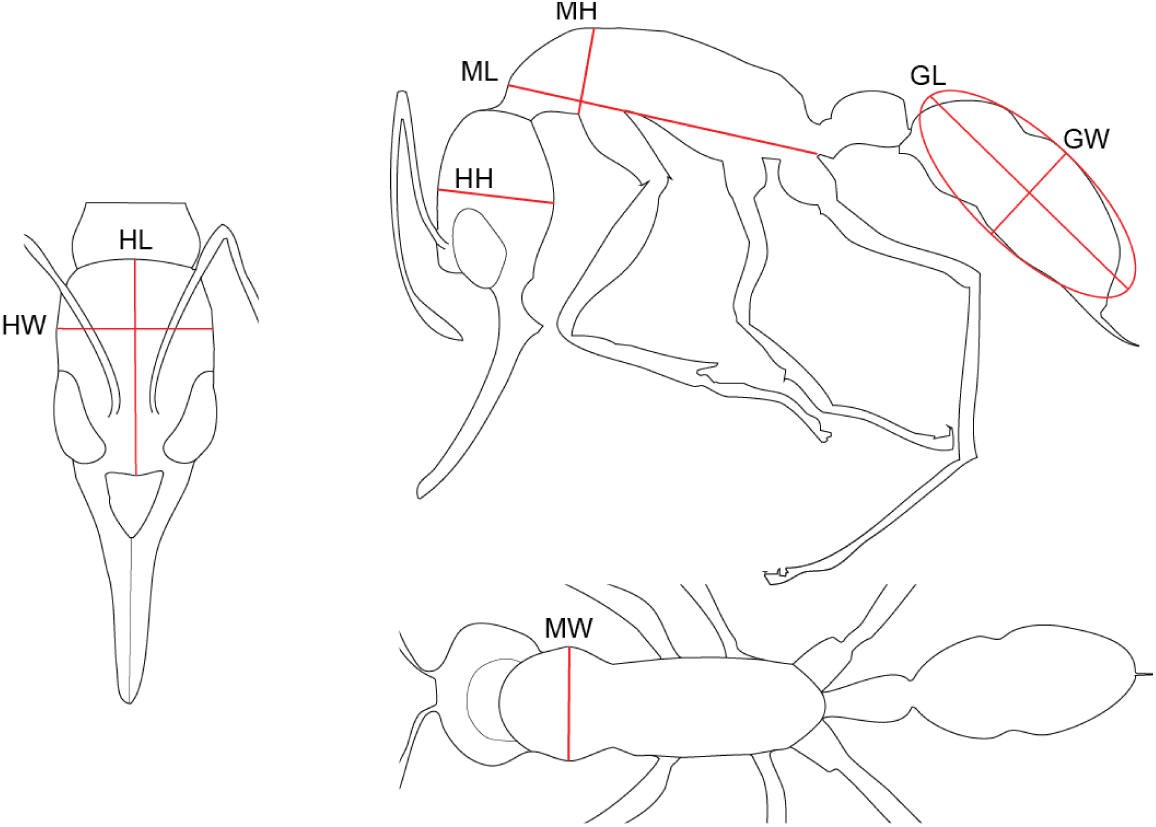
Morphological measurements used in the present study: head high (HH), head length (HL), head width (HW), mesonotum high (MH), mesonotum length (ML), mesonotum width (MW), gaster length (GL), gaster width (GW).

For species with no morphologically distinct reproductive caste (e.g., *Diacamma*), only one individual was measured and categorized both as queen and worker, resulting in a dimorphism value of 1. Species where the measured worker happened to be bigger than the measured queen (e.g., *Mystrium oberthueri* (Molet et al. 2007) and other species with similar sized queens and workers) were also categorized with a size dimorphism value of 1.

Statistical analyses were performed and graphs were generated with the software R (v.4.1.3) (Development Core Team R 2013). All our measurement data are provided in Supplementary material 1. Queen and worker volume values were used as log_10_ transformed and dimorphism ratio were used as log_10_ and root-square transformed in our analyses to fit a normal distribution (Garland Jr et al. 1992, Mongiardino Koch et al. 2015).

### Larval alloparental care and morphologies

Since there is limited information available in the literature about alloparental care and larval feeding across the ant phylogeny, we wanted to examine the relationship between larval feeding habits and their morphologies. Our dataset included information on the larval feeding habits of 99 species spanning 71 genera and 12 subfamilies. Specifically, we categorized whether larvae of these species usually fed semi-autonomously with limited alloparental care (*viz*., actively feeding upon unprepared prey left at their disposal or receiving large chunks of food items) or, conversely, if their feeding primarily involves strong adult care (*viz*., regurgitation, preparation of pre-masticated pellets, distribution of trophic eggs, etc.) (see Supplementary material 1 {data feeding pattern} for quotes and classification).

Over a span of 66 years, Wheeler and Wheeler performed a thorough analysis of larval morphology in more than 700 ant species. Among the global ant larva diversity, they identified 13 general body shapes and 20 general mandible shapes in which they classified ants at the genus level, describing, among other features, the presence or absence of an elongated neck and medial teeth on the mandible (Wheeler and Wheeler 1976, 1986b) (See Fig. S5 and Tables S4-S5 for details about these categories).

According to Wheeler and Wheeler’s statements about the role of neck and mandibles in larval feeding (see results), we retrieved the presence of an elongated neck and medial (sub-apical) teeth on mandibles in larvae to associate these features with their feeding habits. We assembled data on larval morphology for 733 species from 202 genera, mainly from the pioneering work of Wheeler and Wheeler and complemented by some more recent larval descriptions. Other aspects of larvae might also be informative (e.g., mandible size, mandible sclerotization, rate of movement, dynamic range of neck) but there is little to no available data for these traits.

To associate larval morphologies with species from Wheeler and Wheeler’s work, we retrieved morphological categories associated to genera by Wheeler and Wheeler and the list of species they studied (1976, 1986b). We associated species with the morphologies of their genus and updated the names of species according to the latest taxonomic revisions (AntCat.org).

Wheeler and Wheeler described two larval mandible morphology categories as “with or without medial teeth” (Table S5). For the species falling into these categories, we thoroughly examined their detailed larval descriptions to determine the presence or absence of medial teeth in these species. The resulting database and classification details are provided in Supplementary material 1.

These morphological traits display little evolutionary lability (Fig. 2). Within our dataset, among the 113 genera represented by multiple species (with an average of 5.7 species per genus), only five genera displayed intra-genera variability in neck or mandible morphology (*viz*., *Cephalotes*, *Myopias*, *Neivamyrmex*, *Nylanderia*, *Tetramorium*). As such, we matched the species in our dimorphism dataset with larval morphology extrapolated from the genus level, except species belonging to genus with variable larval morphologies that we kept at the species level.

### Social complexity and life history traits

We collected several traits related to the social and ecological evolution of ants. We gathered data on the presence of polymorphic workers at the species level from La Richellière et al. (2022). Polymorphism was considered absent in 11 species from our dataset not included in La Richellière et al. (2022), as polymorphism has been recorded from 0 to 2% in species within their genera. Information on genera with species exhibiting workers with full reproductive potential (gamergates) was retrieved from Antwiki.org. Species in our dataset belonging to these genera were classified as gamergate, and otherwise classified as non-gamergate. We used the classification of Blanchard and Moreau (2017) to retrieve if a genus is mainly predaceous or not. We gathered colony size data from various databases and primary literature for a total of 839 records encompassing 699 species across 184 genera. We calculated the median per species then the median per genus that we extrapolated to the species of our dataset.

### Phylogenetic data

We used the phylogenetic data produced by Economo et al. (2018) as a reference for phylogeny as it was the largest and most recent ant phylogeny data at the time. Specifically, these data include 1) a Maximum Clade Credibility (MCC) backbone tree reconstructed from the sequence alignment of 679 specimens, 2) a set of 200 topologies in which almost all (∼15k) ant species were randomly grafted around the leaves of the same genus from backbone trees, and 3) one MCC ∼15k species grafted tree. Species names were updated according to the latest taxonomic revisions (AntCat.org, release March 2023). Species of our dataset missing in the grafted trees were included at the position of another unused species of the same genus.

### Modeling trait evolution

We investigated the temporal dynamics of morphological evolution over the diversification of ants using the MCC backbone tree, whose leaves are fully supported by molecular data. For each trait (i.e., body volume of queens and workers and queen:worker dimorphism), we assessed their phylogenetic signal in the data by calculating Pagel’s lambda and Blomberg’s *K* with the R package phytools (Revell 2012) (v.1.9.16). We then tested the fit of nine models of continuous trait evolution for each trait: we applied a Brownian motion model, a single-optimum Ornstein–Uhlenbeck model, an early burst model, a white noise model, a rate trend model, a lambda model, a kappa model, and a delta model of trait evolution using the function *fitContinuous* of the R package Geiger (Harmon et al. 2008) (v.2.0.10). The models were compared by their log-likelihood and Akaike information criterion (AIC). Results obtained from trait evolution analyses are summarized in Table S2.

Differences in evolutionary rates (σ^2^) of traits were assessed following Adam’s method (2013) implemented in the package mvMORPH (Clavel et al. 2015) (v.1.1.7). We compared the log-likelihood of two Brownian motion based models, one where the two traits or clades to compare evolve at distinct evolutionary rates, and another where the traits or clades are constrained to evolve at a common evolutionary rate. The log-likelihood values of the two models were compared using the Likelihood Ratio Test hosted in mvMORPH (Clavel et al. 2015) (v.1.1.7).

### Ancestral state reconstructions

Ancestral state reconstructions were performed on the MCC ∼15k species grafted tree. We performed ancestral state reconstructions for larval body shape and mandible shape at each node using the function *fitMk* in the R package phytools (Revell 2012) (v.1.9.16). This utilizes a stochastic character mapping approach (Huelsenbeck et al. 2003, Bollback 2006). We compared three evolutionary models of state transitions (equal-rates, symmetric and all-rates-different) using their Akaike information criterion weight (AICw). Ancestral state reconstructions of the Q:W dimorphism were performed using the function *anc.ML* in the R package phytools (Revell 2012) (v.1.9.16), where we estimated evolutionary parameters and ancestral states using likelihood given a Brownian-motion model of evolution of continuous traits. To incorporate the best-fitted lambda evolutionary model and account for the phylogenetic signal of dimorphism (as detailed above), ancestral state estimations were performed on a lambda-transformed tree. The tree was transformed according to the calculated phylogenetic signal of dimorphism using the function *rescale* of the R package Geiger (Harmon et al. 2008) (v.2.0.10). We then projected reconstructed ancestral states onto the original species tree.

### Correlation between traits

We tested the evolutionary correlation between larval feeding pattern and larval morphology features (i.e., presence of an elongated neck and medial teeth) on the MCC ∼15k species grafted tree using the function *fitPagel* hosted in the R package phytools (Revell 2012) (v.1.9.16). A function designed to fit Pagel’s (1994) method testing for an evolutionary relationship between two binary characters. The evolutionary relationships between traits were investigated by building additional models where one trait depends on the other. The model fits were then compared by their Akaike information criterion (AIC) (Table S1).

We used Phylogenetic Generalized Least Squares (PGLS) analysis to test the correlation between traits taking into account relatedness among taxa. PGLS uses a covariance matrix to weigh least squares and incorporate phylogenetic relatedness into the analysis, assuming that branch length is proportional to the residual error in the model (Felsenstein 1985, Revell 2010). As such, we performed a PGLS analysis using the function *gls* of the R package nlme (Pinheiro et al. 2018) (v.3.1.157) on the mean covariance matrix between species from the 200 phylogenetic trees. Before calculating the covariance matrix, trees were lambda transformed, using the function *rescale* of the R package Geiger (Harmon et al. 2008) (v.2.0.10), according to the phylogenetic signal of the tested answer variable (see Modeling trait evolution), a classical method to implement phylogenetic signal in PGLS analysis (Symonds and Blomberg 2014). Log-transformed dimorphism values were additionally root-squared transformed in order that residuals fit a normal distribution.

To refine our understanding of how traits influenced dimorphism, we used phylogenetic path analysis (von Hardenberg and Gonzalez-Voyer 2013) implemented in the R package Phylopath (van der Bijl 2018) (v.1.1.3). This method allows the comparison of models of possible causal relationships between traits while testing for direct or indirect effects using the d-separation method (Shipley 2000, 2009) and considering the non-independence of the traits due to phylogeny through phylogenetic generalized least squares analysis (PGLS). We built several plausible models and estimated their parameters on the MCC tree. Following van der Bijl (2018), we selected models with the minimum corrected Akaike Information Criterion (CICc) within a range of 2 CICc and with no significant p-values indicating conditional independencies. To account for uncertainties in our phylogenetic data, we also retrieved the model selection over the 200 grafted trees to corroborate results from the MCC tree. Additionally, we edited the function *average()* from the phylopath package to handle a set of trees instead of a single one, and estimate coefficients of variables’ relationships arising from model selection over the 200 grafted trees and average them using the function *average_DAGs()*. We provide all our R codes to conduct these analyses in the supplementary files.

## ACKNOWLEDGMENTS

We would like to dedicate our study to Jeanette and George Wheeler, whose entire carrer focused on the description of ant larvae permited this analysis. We are grateful to the contributors and curators of AntWeb and AntCat for making freely available to the public a vast store of images and information about ants. We thank Fuminori Ito, Daniele Silvestro, Sanja Hakala and Marie-Pierre Meurville for their feedback on our manuscript. We acknowledge Riou Mizuno for valuable discussions about ant evolution, François Brassard for providing us with pictures of ants, and Wouter van de Bijl for his help with phylopath. A.C.L. was supported by the Swiss National Science Foundation (PR00P3_179776).

## SUPPLEMENTARY TABLES AND FIGURES

**Table S1.** Coevolution model comparisons for larval feeding habits, larval morphology, and species diet. Results from Pagel’s (1994) method for testing for evolutionary relationships between binary characters on a dataset of 99 species (71 genera) shown in Figure S1.

| y \ x | Supervised larval feeding habits |  |  |  |  |
| --- | --- | --- | --- | --- | --- |
|  | model | d.f. | log(L) | AIC | weight |
| Neckless larvae | independent | 4 | -81.79004 | 171.5801 | 7.9438E-06 |
|  | interdependent x & y | 8 | -66.10942 | 148.2188 | 0.9394206 |
|  | x dependent of y | 6 | -73.88248 | 159.765 | 0.00292179 |
|  | y dependent of x | 6 | -70.90029 | 153.8006 | 0.05764969 |
| Toothless larvae | independent | 4 | -81.59694 | 171.1939 | 9.3427E-06 |
|  | interdependent x & y | 8 | -67.25114 | 150.5023 | 0.2907999 |
|  | x dependent of y | 6 | -70.91164 | 153.8233 | 0.05526485 |
|  | y dependent of x | 6 | -68.44078 | 148.8816 | 0.6539259 |
| Non-predatory diet | independent | 4 | -84.89965 | 177.7993 | 0.00111907 |
|  | interdependent x & y | 8 | -75.35664 | 166.7133 | 0.2858611 |
|  | x dependent of y | 6 | -76.4453 | 164.8906 | 0.7111222 |
|  | y dependent of x | 6 | -82.3715 | 176.743 | 0.00189771 |

**Table S2.** Comparison of model fits for different models of trait evolution and phylogenetic signal for body volumes and dimorphism. Models were estimated with a subset of 103 species on a tree whose phylogenetic relationships are fully supported by molecular data.

| Trait | Model comparison<br>log-likelihood difference (AIC difference) |  |  |  |  |  |  |  | Phylogenetic signal |  |  |  |
| --- | --- | --- | --- | --- | --- | --- | --- | --- | --- | --- | --- | --- |
| | Brownian motion | Ornstein-Uhlenbeck | early busrt | white noise | rate trend | mean trend | lambda | kappa | delta | $\lambda$ | K | $\sigma^2$ |
| Queen volume | -5(8) | 0(1) | -5(10) | -7(11) | -1(3) | -5(10) | -3(6) | -5(9) | **0(0)** | 0.867 | 0.56 | 0.0093 |
| Worker volume | -2(2) | **0(0)** | -2(4) | -19(36) | 0(0) | -2(4) | -1(3) | -2(4) | 0(0) | 0.894 | 0.77 | 0.0058 |
| Q:W dimorphism | -4(6) | -1(2) | -4(8) | -8(15) | -1(3) | -4(8) | **0(0)** | -1(2) | -1(2) | 0.847 | 0.61 | 0.0034 |

**Table S3.** Details of values from PGLS analysis conducted on dimorphism, queen volume and worker volume on 363 species over 128 genera (df = 357). Dimorphism values were log_10_ and root-squared transformed in order that residuals fit normal distribution according to the Shapiro test.

| Trait | sqrt(log(Q:W dimorphism))<br>$\lambda = 0.847$ | | | | log(queen volume)<br>$\lambda = 0.867$ | | | | log(worker volume)<br>$\lambda = 0.894$ | | | |
| --- | --- | --- | --- | --- | --- | --- | --- | --- | --- | --- | --- | --- |
|  | Value | Std.Error | t-value | p-value | Value | Std.Error | t-value | p-value | Value | Std.Error | t-value | p-value |
| larval passiveness | 0.121 | 0.034 | 3.522 | **0.00048** | -0.078 | 0.084 | -0.925 | 0.35559 | -0.267 | 0.079 | -3.369 | **0.00084** |
| predatory diet | 0.024 | 0.045 | 0.544 | 0.58675 | -0.119 | 0.109 | -1.093 | 0.27498 | -0.205 | 0.103 | -1.993 | 0.04705 |
| log colony size | 0.055 | 0.014 | 3.856 | **0.00014** | 0.242 | 0.035 | 6.939 | **1.9E-11** | 0.133 | 0.033 | 4.065 | **5.9E-05** |
| worker polymorphism | 0.180 | 0.033 | 5.502 | **7.2E-08** | 0.533 | 0.080 | 6.676 | **9.4E-11** | 0.277 | 0.075 | 3.719 | **0.00023** |
| reproductive workers | -0.166 | 0.069 | -2.424 | **0.01584** | 0.512 | 0.168 | 3.046 | **0.00249** | 0.584 | 0.158 | 3.691 | **0.00026** |

**Table S4.**
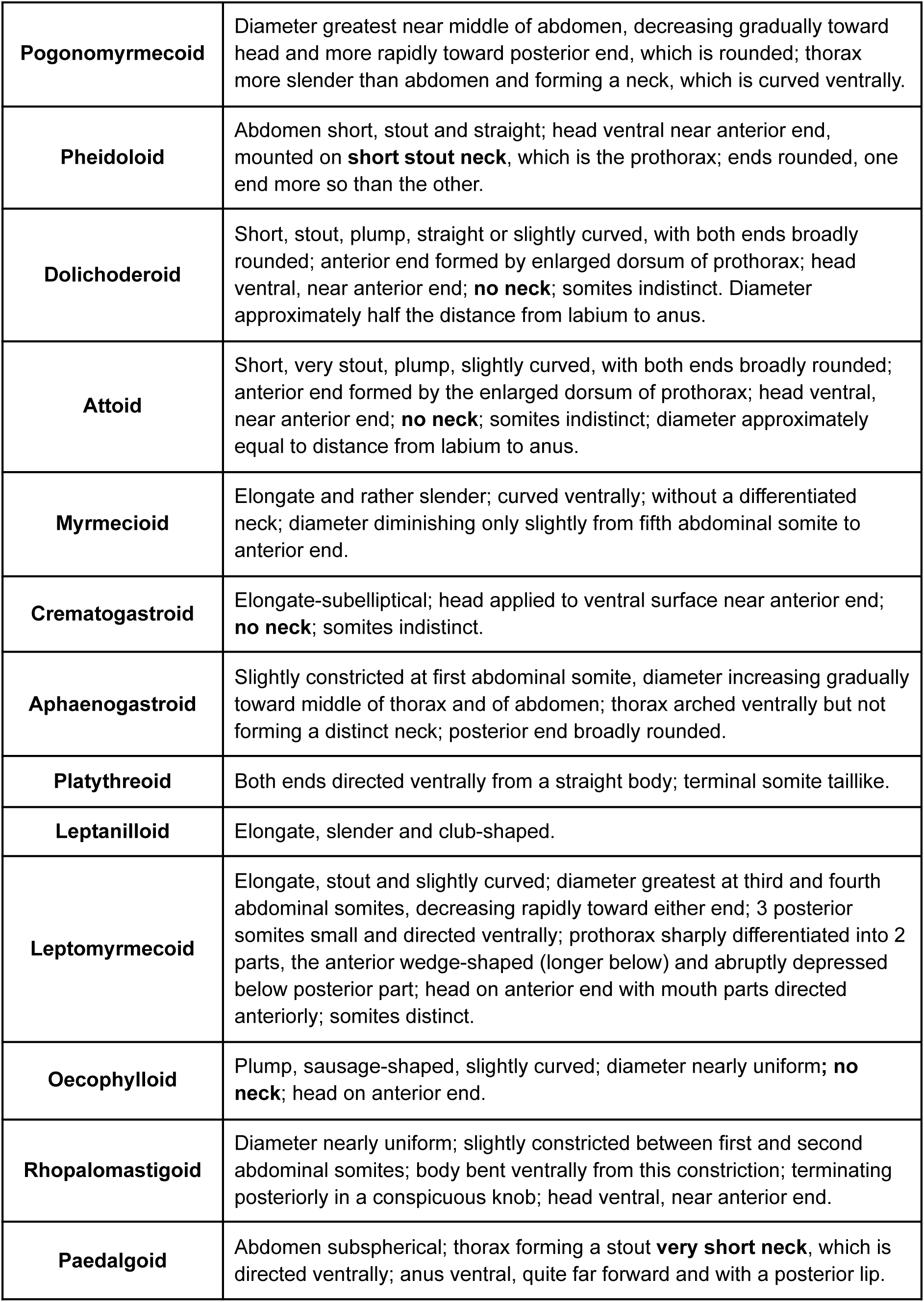
Descriptions of the ant larvae’ body shape categories provided by Wheeler and Wheeler (1976, 1986). Are in bold the elements indicating larval passivity.

**Table S5.**
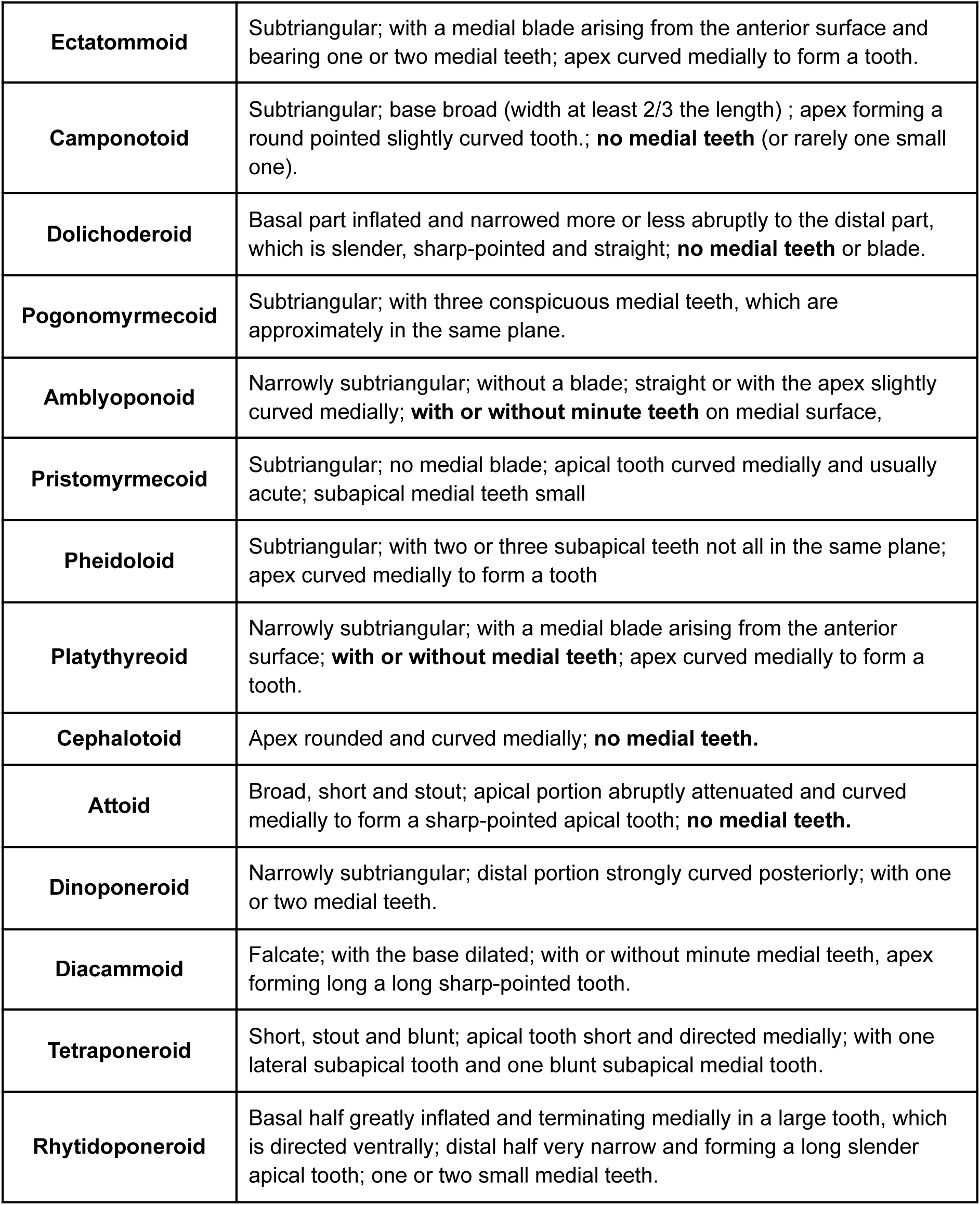
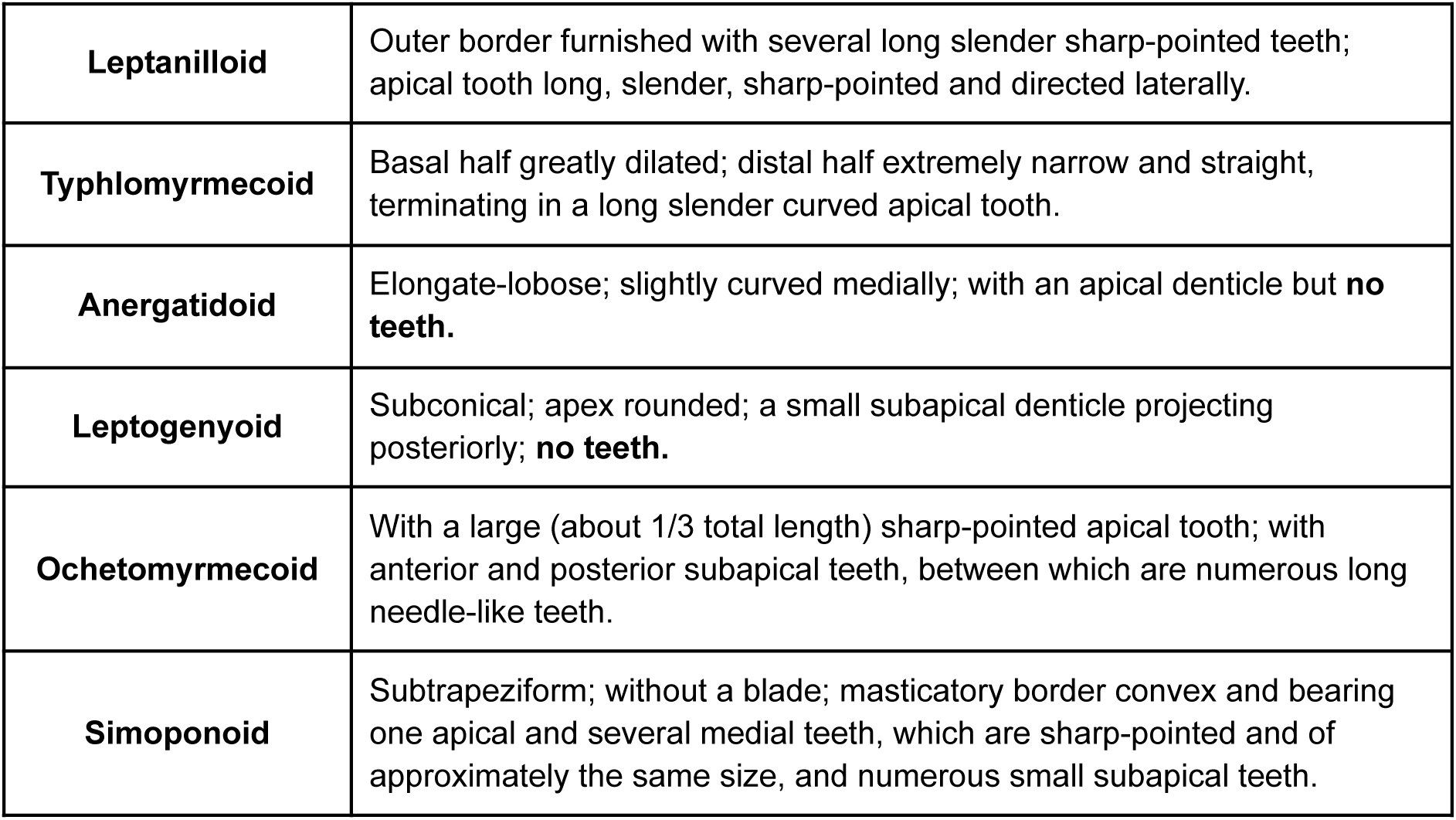
Descriptions of the ant larvae’ mandible shape categories provided by Wheeler and Wheeler (1976, 1986). Are in bold the elements indicating larval passivity. For species falling into categories described as “with or without medial teeth” we examined their detailed larval descriptions to determine the presence or absence of medial teeth in these species.

**Figure S1.**
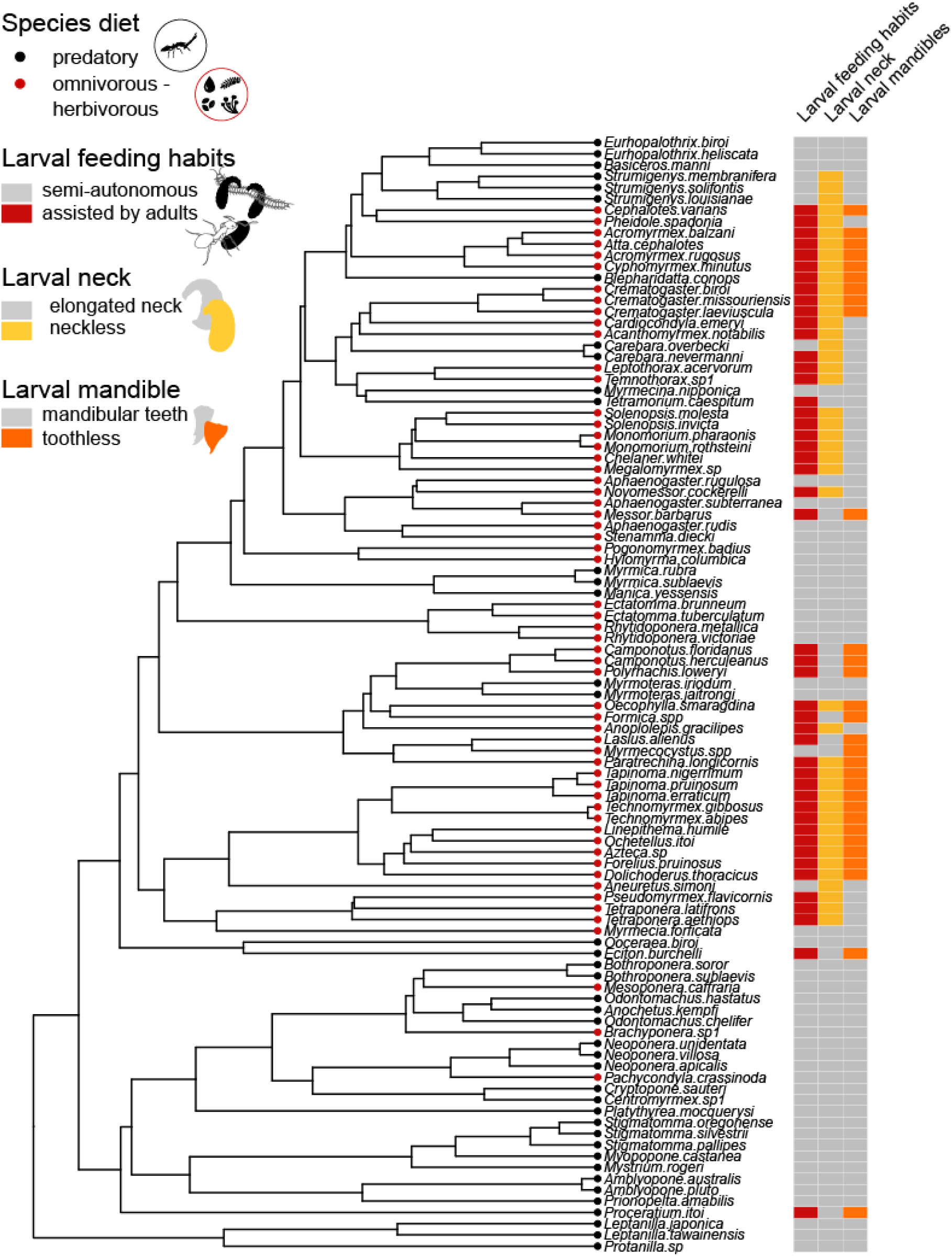
Phylogenetic representation of the 99 species covering 71 genera with both data for larval feeding habits, larval morphology, and species diet.

**Figure S2.**
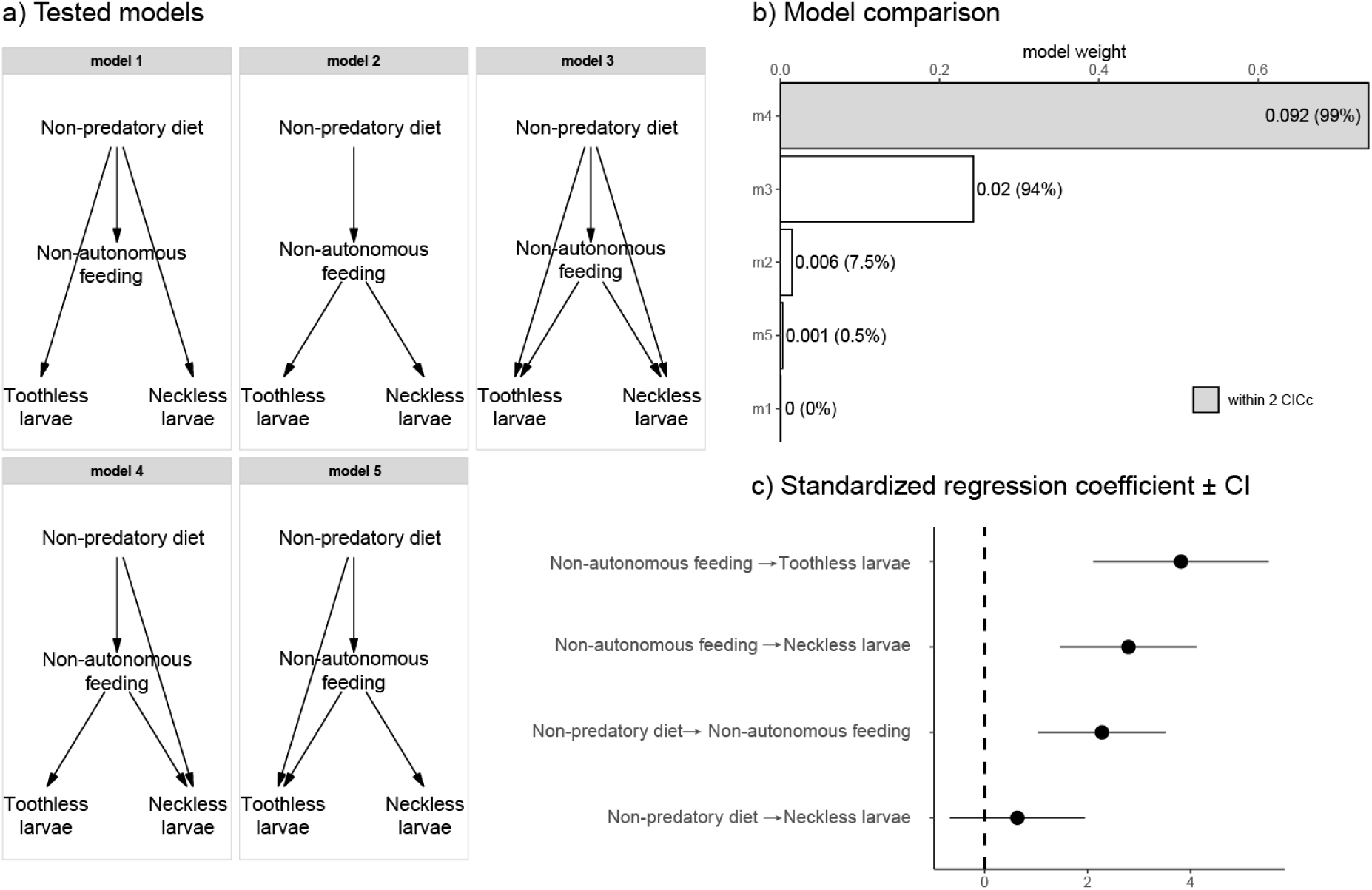
Supplementary information for Figure 2. Multi-tree phylogenetic path analysis performed on the dataset of 99 species (71 genera) shown in Figure S1. **a**) Directed acyclic diagrams of the compared models. **b**) Model comparison and selection (in grey). Bar labels are p−values, significance indicates model rejection; values between parentheses are the percentages of model selection over 200 phylogenetic trees. **c**) Standardized regression coefficient ± CI estimated over 200 phylogenetic trees from the selected model (model 4).

**Figure S3.**
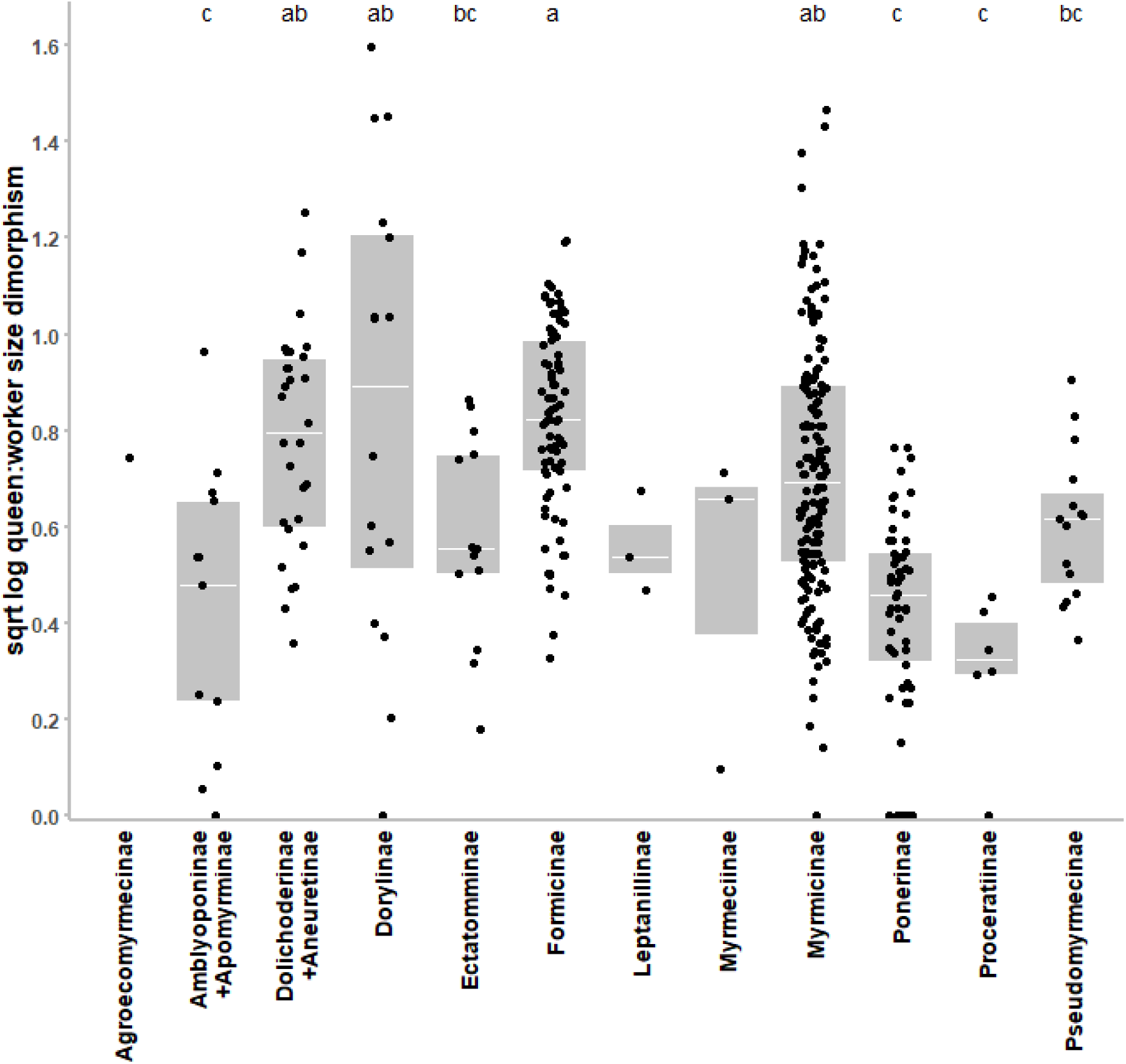
Dimorphism values measured in 392 species distributed in the different ant subfamilies. Ant subfamilies differ in their dimorphism values (one-way ANOVA: F_11,376_ = 15.812, p < 0.001). Different letters indicate significant differences between subfamilies (Tukey’s post-hoc test: p < 0.05). Groups containing less than 6 species were not considered in the statistical analysis. Dimorphism values were log_10_ and root-squared transformed in order that residuals fit normal distribution according to the Shapiro test.

**Figure S4.**
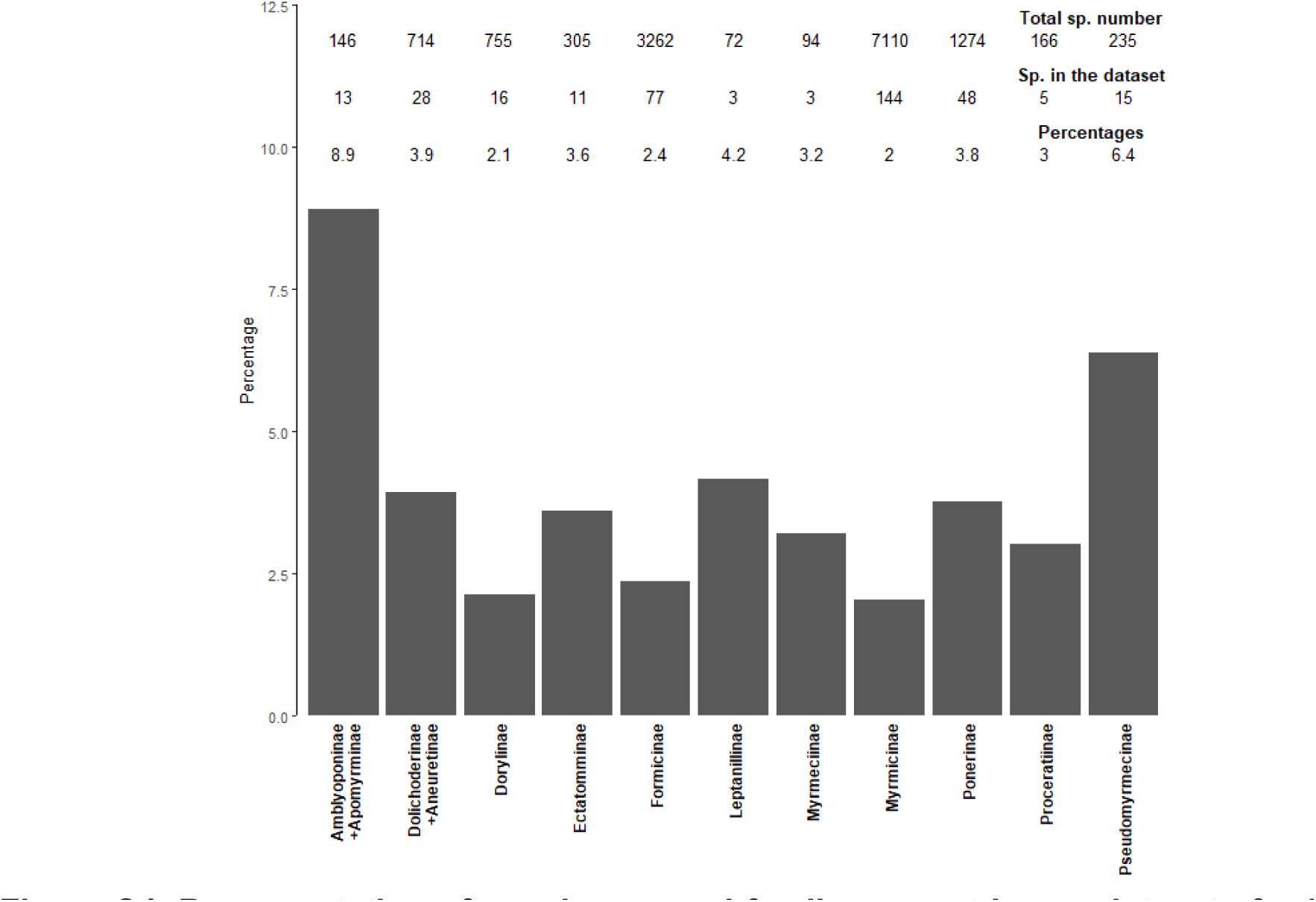
Representation of species per subfamily present in our datasets for PGLS analysis. Total number of species data is from Antcat.org (September 2023).

**Figure S5.**
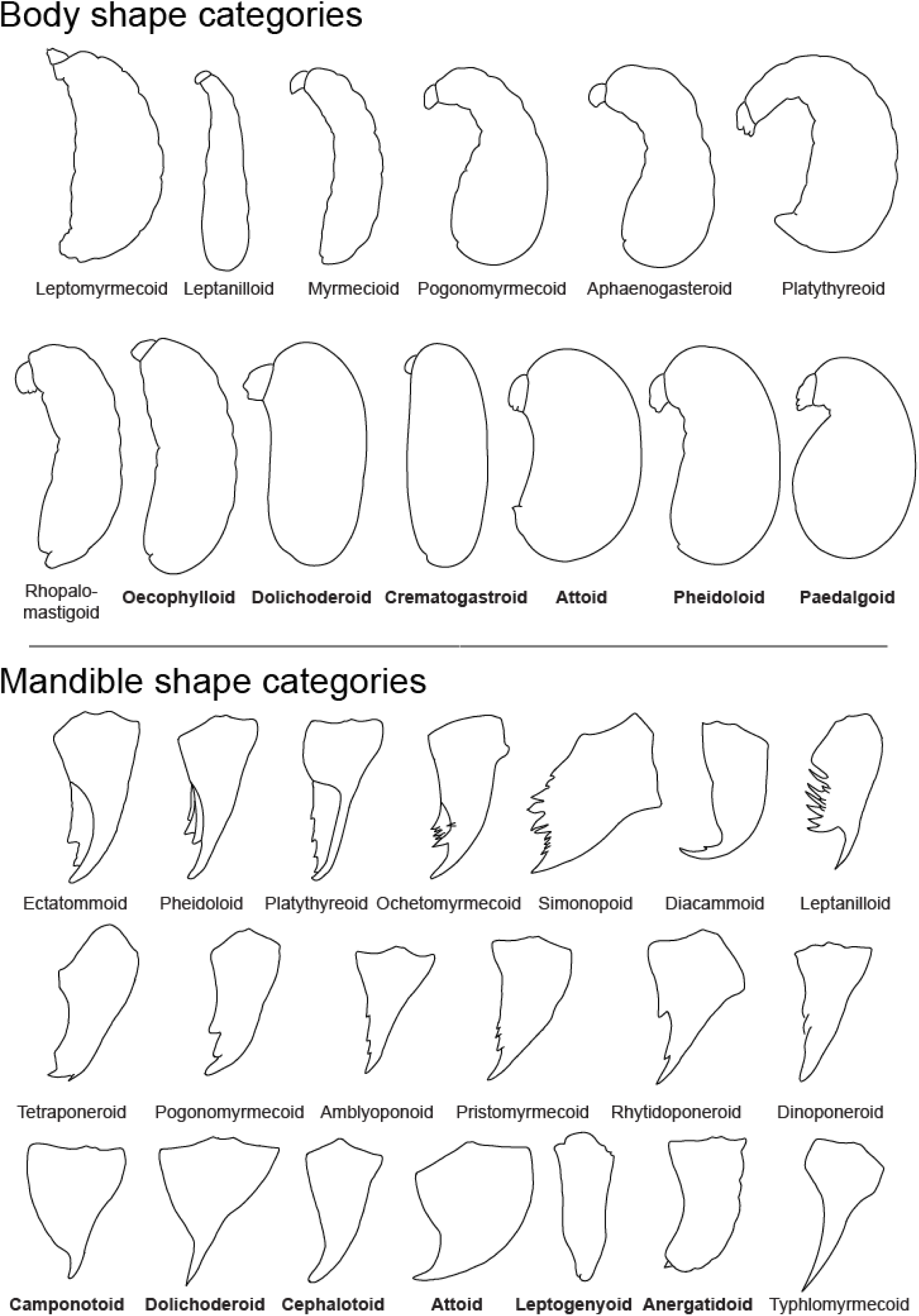
Ant larval body and mandible shape categories proposed by Wheeler and Wheeler (1976, 1986). Names in bold are categories considered ‘passive’ according to the absence of elongated neck and absence of mandibular teeth following their description (see Tables S4 and S5). Although the Typhlomyrmecoid mandible category appears toothless in the Wheeler and Wheeler representation shown, the description of the category does not mention the absence of medial teeth and the larvae of the concerned genera (*Apomyrma* and *Typhlomyrmex*) are described and represented with medial teeth (Wheeler and Wheeler 1964, Theeler and Wheeler 1970).

## SUPPLEMENTARY DISCUSSION

### Changing larval diet requires other changes

Transitioning to alternative food sources is a complex process. Since ant larvae arose from a prey-feeding background, any new food still must fulfill nutritional requirements for larval growth; i.e., rich in protein and nitrogen (Dussutour and Simpson 2009, Csata and Dussutour 2019). Different ant lineages overcame this challenge through diverse innovations. For instance, sugary liquids can be enriched by nitrogen-fixing bacterial symbionts inside the digestive tracts of workers (Borm et al. 2002, Russell et al. 2009, Jackson et al. 2022), or through metabolic labor performed by workers (Hakala et al. 2021, Negroni and LeBoeuf 2023b). Both innovations entail worker ingestion, followed by sharing through trophic eggs or regurgitation, two features common in species with passive larvae. Other ants, including fungus-growing ants and some Pseudomyrmecinae, forged relationships with external symbionts such as fungi or plants that provided nutritional support for larval growth (Janzen 1966, Quinlan and Cherrett 1979, Blatrix et al. 2012).

### Behavioral and temporal innovations strengthening adult control over larvae

Additional behavioral adaptations likely contributed to the evolution of passive larvae. Age-related changes in larval diet have been reported in several species, with young larvae receiving regurgitate while late-stage larvae are provided with prey (e.g., Petralia and Vinson 1978, Buschinger and Schaefer 2006). Besides the potential limited ability of young larvae to handle prey, the pressure to monitor larval feeding may also decrease as their developmental trajectory settles (Abouheif 2021). Turning prey into pellets instead of leaving larvae consuming it directly ensures individualized and monitored feeding, albeit placing a greater burden on workers. These innovations were likely largely facilitated by the use of alternative food sources for larvae, reducing the amount of prey to process and associated with larger colony sizes (Meurville et al. 2022). Larger colonies often have greater task organization (Thomas and Elgar 2003), allowing the evolution of other facilitators such as sorting larvae by stage (Franks and Sendova-Franks 1992) and nurse larval-stage specialization (Walsh et al. 2018), which likely strengthen the regulation of larval growth. In addition, forming cohorts of larvae that become either all queens or all workers may be another way for adults to efficiently manipulate larvae’ caste fate. By separating larvae spatially within the nest or temporally (e.g., seasonality), the colony can overcome the challenge of having to create inequality among larvae with similar caste fates. However, the sporadic recording of these behaviors precludes macroevolution analysis at this time.

### Several remaining mysteries on the evolution pattern of passive larvae

Regarding these associations between species diets, larval feeding habits, larval morphology and size dimorphism, several questions remain unanswered on the evolution pattern of passive larvae. Regarding the consistent evolution of toothless larvae in dorylinae Army-ants, Wheeler and Wheeler pondered in 1984 “*Why do larvae of the world’s most carnivorous ants have mandibles incapable of chewing solid food?*”. Although Wheeler and Wheeler (1984) speculated that these larvae might digest prey externally before sucking the resulting liquid, the mystery remains unsolved, and the cause and consequence of such mouthpart reduction are even more obscure. This particular adaptation may arise from their diet largely composed of other ants’ brood (Hoenle et al. 2024). On the other hand, several species of *Leptogenys* and *Myopias* have also evolved toothless morphologies in otherwise actively feeding larvae. The case of these species, however, might stem from specialization on prey with extremely thick cuticles, such as millipedes, which restricts larvae to feed only on the internal soft tissues, as described by Brown (1992) and Ito et al. (Ito et al. 2020). Finally, it is interesting to note that many species retain elongated necks in lineages that have first evolved toothless larvae (e.g., Formicinae) While the necessity for mandibular teeth may primarily relate to food texture handled by larvae, the pressures influencing larval necks can be considerably more diverse. Larval opportunities for enhanced mobility, food-begging behaviors, brood cannibalism, and handling larger prey may favor the presence of necks in larvae (Urbani 1991, Peignier et al. 2019). Additionally, the already-increased worker control over larval feeding in those lineages may alleviate pressure for entirely passive larvae.

